# High occurrence of plasmid-mediated quinolone and ESBL resistance genes among multidrug resistant *Escherichia coli* from clinical samples in two healthcare facilities in Yaounde, Cameroon

**DOI:** 10.64898/2026.08.07.743442

**Authors:** Priscille Koubissak Mbende, Jaures Kenfack Noumedem, Luria Leslie Founou, Ange Atsafack Zobou, Jannet-Vanèle Meli, Raspail Carrel Founou

## Abstract

**Introduction:** In sub-Saharan Africa, and more specifically in Cameroon, antimicrobial resistance (AMR) represents a major public health threat. This is underlined by the increasing appearance of multidrug-resistant bacteria. Extended-spectrum β-lactamase producing *Escherichia coli* (ESBL-Ec), a critical priority bacterium, is increasingly implicated in life-threatening infections in hospital and community settings in Cameroon. Data on the genetic composition of ciprofloxacin-resistant *Escherichia coli* are limited in Cameroon. This study aimed to investigate the prevalence, genetic diversity, resistance mechanisms in multidrug-resistant *Escherichia coli* organisms isolated from clinical samples in two hospitals in Yaoundé, Cameroon.

**Method:** A cross-sectional study was conducted from February to June 2025 in two healthcare facilities in Yaounde, Cameroon. All clinical samples from in- and out-patients were analysed. After culturing, identification was performed using API20E as per the manufacturer’s instructions and ESBL production was screened in CHROMagar^TM^ ESBL. Antimicrobial susceptibility testing was performed using the Kirby-Bauer disc diffusion method. Polymerase chain reaction (PCR) was used to detect ESBL and plasmid mediated quinolone resistance (PMQR)genes, as well as mutations in quinolone resistance-determining region (QRDR) (*gyrA*/*parC*) Horizontal. plasmid transfer was also investigated. Finally, phylogroup analysis was assessed.

**Result:** The prevalence of MDR *E. coli* was 50.7% (n=33/65), all of which (100%) were ESBL producers and 91% were ciprofloxacin-resistant. Highest resistance rates were observed for cefotaxime (100%), ceftriaxone (100%), and ciprofloxacin (91%). The most frequent ESBL genes were *bla*_TEM_ (36.3%; n=12/33). Among PMQR genes, *qnrB* was detected in 16.6% (n=5/30) of isolates. Only the ESBL genes were carried by plasmids; the most prevalent plasmid-borne gene was *bla*_TEM_ (40%), followed by *bla*_CTX-M_ (26.7%). Mutations within the topoisomerase QRDR (parC gene) were identified in 36.6% (n=11/30) of ciprofloxacin-resistant strains. Phylogroup analysis revealed a predominance of phylogroup A, followed by group B.

**Conclusion:** This study reveals a high prevalence of multidrug-resistance, ESBL (*bla*_TEM_ *dominant)* and fluoroquinolone resistance in *E. coli* in Yaoundé, with plasmid dissemination of ESBL genes and chromosomal stabilization of PMQR determinants. The predominance of commensal phylogroups in clinical samples underlines the role of the community reservoir. It is urgent to reinforce " real-time One Health" genomic surveillance in Cameroon.

## INTRODUCTION

Antimicrobial resistance (AMR) remains a global public health problem, especially in low-and-middle income countries (LMICs), like Cameroon, where lack of water, sanitation and hygiene (WASH) combined with suboptimal infection prevention and control measures (IPC) in healthcare facilities aggravate this challenge (1). Bacterial AMR was the leading cause of mortality in 2019 associated with 4,5 million deaths globally and will reach 10 million deaths by 2050 if nothing is done to contain it (2). During the high-level meeting of the United Nations General Assembly (UNGA) in 2024, world leaders pledged 10% reduction of AMR-related mortality by 2030 (3). This Jeddah ministerial conference on antimicrobial resistance reaffirmed the urgency of a coordinated One Health response. Its commitments aim to strengthen governance, financing, surveillance, rational antimicrobial use, equitable access to treatments, prevention, research and international political follow up by 2030 through an integrated, multisectoral approach across human, animal, and environmental sectors (3).

ESBL-producing *Enterobacterales* are classified as critical priority pathogens by the World Health Organisation (WHO),owing to their high burden of disease (mortality and morbidity), increasing resistance trends, and the scarcity of promising new antibiotics (4). Among these, ESBL-producing *Escherichia coli (ESBL-Ec)* is of particular concern because of its limited treatment options, widespread dissemination and rapid development of resistance to broad-spectrum antibiotics, including β-lactams and fluoroquinolones. Additionally, infections caused by multidrug-resistant *Escherichia coli* (MDR-Ec) are associated with a substantial clinical and public health burden, including increased treatment failure, prolonged hospitalization, and higher mortality rates(5). *E coli* represents one of the most important pathogens associated with healthcare-acquired infections (HAIs) including blood stream infections (BSIs) and urinary tract infections (UTIs) in Cameroon (6) (7).

In Africa particularly in Cameroon, clinical *E. coli* isolates often show high multidrug resistance (8, 9) (10). The accessibility and availability of fluoroquinolones in hospital and community settings led to the emergence of fluoroquinolones resistance among ESBL-*Ec* in many sub-Saharan African countries like Cameroon; Nigeria, Egypt, South Africa(10, 11).

Fluoroquinolones (FQs) are broad-spectrum antibiotics, commonly used in sub-Saharan Africa, particularly in Cameroon, as effective antibiotics for the treatment of infectious diseases caused by ESBL-E*. coli* in hospitals (10, 11). Unfortunately, in order to evade the activities of fluoroquinolones, these bacteria have developed resistance mechanisms, including chromosomal mutations in the determining region of quinolone resistance (QRDR) via DNA gyrase and topoisomerase enzymes and plasmid-mediated quinolone resistance (PMQR) including the *qnr* genes family(*qnrA, qnrB, qnrC, qnrD and qnrVC21)* (11, 12) (13). Finally, *qepA* and *oqxAB* genes which enhance efflux pump activities, reduce bacterial susceptibility to quinolones and also amplify extended-spectrum ß-lactamase activity,thereby contributing to the emergence and further complicating clinical management (11, 12).

Futhermore, understanding the phylogroup of *E. coli* provides useful insight into the distribution of resistance and virulence traits across strains, and can help identify lineages more frequently associated with clinical infection versus commensal carriage. Based on the classical Clermont phylogroup scheme, *E.coli* is classified into seven phylogroups (A, B1, B2, C, D, E and F) (14) with strains involved in clinical infections often associated with group B2 and, to a lesser extent, group D, while groups A and B1 are more frequently associated with commensal and less virulent (14).

In Cameroon, several recent studies have documented a high prevalence of MDR and ESBL-producing *E. coli* across different setting and regions. In the West region, MDR and *ESBL-Ec/Kp* were highly prevalent among urinary tract infections(6). In Yaoundé, *ESBL-Ec* was detected in the majority of pigs and exposed slaughterhouse workers, illustrating the one Health dimension of transmission(15). More recently, the first report of report of PMRQ among *ESBL-Ec/Kp* in Cameroon was documented among hospitalized patients, hospitalized surfaces, and wastewater in two healthcare setting of Yaounde (16). A parallel phenotypic and genotypic characterization of PMQR and ESBL-producing *Enterobacterales* was recently published in two healthcare facilities in Douala confirming the high prevalence of *E.coli* follow by *Klebsiella pneumoniae* in clinical samples (17). However, comprehensive data combining phenotypic and genotypic characterization including phylogenetic group distribution among *MDR-Ec* remain scare in the country. Furthermore, the extent to which chromosomal QRDR mutations versus plasmid-mediated mechanisms drive fluoroquinolone resistance in this setting is still poorly characterized.

This study therefore aims to investigate the prevalence, phenotypic profiles and genetic characteristics including, plasmid borne antibiotic resistance genes and point mutation regions conferring resistance to fluoroquinolones among *MDR-Ec isolated* from clinical samples in two hospitals in Yaounde, Cameroon.

## MATERIAL AND METHODS

### 1. Study design, setting and period

This study was cross-sectional and conducted during five months, from February to June 2025 at the Yaounde Central Hospital (YCH) and the Yaounde University Teaching Hospital (YUTH), which are tertiary-categories hospitals. The YCH is one of the largest health facilities in Yaoundé, the capital city of Cameroon. It provides patients with internal and specialty medicine units, surgery, emergency room, intensive care units, outpatient department, obstetrics and gynaecology department, pharmacy, laboratory, radiology and pathology. YUTH takes care of many patients requiring prolonged hospitalization and also trains medical professional in the country’s health system. It includes an emergency department, an obstetrics and gynaecology department, a paediatric department, an internal medicine department, intensive care, a laboratory, an outpatient department, a mortuary department, a pharmacy department and a hygiene and maintenance department.

### 2. Sample collection, culture and identification

A total of 754 samples were routinely processed in the microbiology laboratories of the two healthcare facilities following standard microbiological techniques during the study period. Clinical samples were composed of urine, sputum, endocervical swabs, faecal samples, cerebro-spinal fluids, blood samples, wound swabs, and catheters. After sampling, all clinical samples were cultured on MacConkey agar and Chromogenic agar CHROMagar^TM^ (CHROMagar, Paris, France) at 37 °C for 24 hours. All grown colonies were identified using API20E (BioMérieux, Marcy l’Etoile, France) following the manufacturer’s instructions. All putative *E. coli* were collected and conserved in cryovials containing trypticase soya broth (Oxoid LTD, Bagingstoke, Hamsphire, England) supplemented with 20% (v/v) glycerol and transported within four hours to the Research Institute of the Centre of Expertise and Biological Diagnostic of Cameroon (<u>CEDBCAM-RI)</u> for ESBL screening and molecular characterization.

### 3. Ethical consideration

To perform this study, four approvals were obtained from the regional Ethics Committee for Human Health Research (Ref: 0086-3/CRERSHC/2025), the Yaounde Central Hospital (N°017/25/AR/MINSANTE/SG/DHCY/CM/), the Yaoundé University Hospital (Ref: 00205-25/QAR/CHUY/DG/DGA/DM/CAPRC/SDSITMS/CEAAP/CEARC/CB), and the Research Institute of CEDBCAM (Ref:2025/02/005/L/CEDBCAM-RI/DG/DRD).

Written informed consent was not obtained because the researchers did not have direct contact with the patients. This study was conducted using anonymized, coded clinical specimens routinely received by the laboratory, and no patient-identifiable information was accessible to the research team.

### 4. Antimicrobial susceptibility testing (AST)

Antibiotic susceptibility testing was performed by a modified Kirby-Bauer disc diffusion method as recommended by the Antibiogram Committee of the French Society of Microbiology of 2024(18). The fresh and pure colonies were used to make an inoculum that was standardized using the 0.5 McFarland control. This inoculum was inoculated on Muller Hinton agar and incubated at 37°C for 24 hours. A panel of 11 antibiotics of the Italian brand Liofilchem belonging to five different classes, including: amoxicillin-clavulanic acid (20/10μg), ceftriaxone (5μg), cefotaxime (30μg), ceftazidime (10μg), meropenem (10μg), ciprofloxacin (5μg), chloramphenicol (30μg), gentamicin (10μg), levofloxacin (10µg), norfloxacin (10μg) and Trimethoprim-sulfamethoxazole (25µg). Any isolate showing resistance to at least three or more different classes of antibiotics was considered MDR(19)

All *E. coli* isolates were screened for ESBL production using chromogenic medium CHROMagar ESBL^TM^ (CHROMagar, Paris, France) and confirmed using the double-disk synergy testing as described previously(18). After screening, all confirmed isolates were purified and stored.

### 5. Genotypic characterization

#### a. Detection of ESBL, PMQR and points mutations resistance genes

Genomic DNA (gDNA) and plasmid DNA (pDNA) extractions of ESBL-*Ec* and ciprofloxacin resistant ESBL-*Ec* isolates were performed using a modified boiling method (15) and the PureLink™ Quick Plasmid Miniprep Kit (Thermo Fisher Scientific, Invitrogen, Paris, France), respectively.

Polymerase chain reaction was performed to identify the genes encoding for resistance to fluoroquinolones (*qnr*B, *qnr*S, *qnr*A) (20), mutation genes (*gyr*A and *par*C)(12), β-lactams (*bla*_TEM_, *bla*_CTX-M_, and *bla*_SHV_) (20) Phylogroup assignement was determined by detecting *CHUA, YjaA, TspE4C2* and *ACEK/ArpA*, by multiplex PCR and *ArpAgpE, and trpAGpC1* by singleplex PCR (Phylogroup genes)(21, 22).

All singleplex PCRs (for the detection of *bla_SHV_, oqxA*, *oqxB*, *ArpagpE, TrpAGpC1)* and *multiplex PCRs (*for the detection of *bla*_CTX-M_ and *bla*_TEM_ genes; qnrA*, qnrB and qnrS, gyrA* and *ParC; chuA, YjaA, TspE4C2 and ACEK/ArpA*) were performed, with a reaction mixture of total volume 10 µL. The reaction mixture consisting of 5 μL of One Taq quick load 2X, 0.1 μL of each primer of concentration 10 μM (Table sup I), 2μL of template DNA, and nuclease-free water. PCR products were revealed after electrophoresis on 1.5% (w/v) agarose gel using 0,5μg/mL ethidium bromide solution for 25 minutes and viewed on a G-BOX.

### 6. Plasmid conjugation assays

In order to evaluate the transfer frequency of the *bla*_CTX-M_ and *bla*_TEM_, donor strains (rifampicin-resistant *E. coli* carrying *the bla*_CTX-M_ and *bla*_TEM_ gene) and recipients (*E. coli* ATCC 45266 susceptible to rifampicin and not ESBL) were brought into contact in a Luria Bertani broth (Thermo Fisher Scientific, Invitrogen, Paris, France) and in order to obtain an optical density of about 0.6 to 600 nm and then incubated at 37°C for 4 h. Equivalent volumes of recipient and donor strains were introduced into 500 μl of Luria Bertani broth and then incubated for 24 hours at 37°C. The mixture was centrifuged at 1000 g for 3 minutes. The resulting pellet was cultured on Luria Bertani agar (Thermo Fisher Scientific, Invitrogen, Paris, France) with supplemented Cefotaxim and rifampicin as previously described (24). To determine plasmid transferability, carrying the *bla*_CTX-M_ and *bla*_TEM_ genes, and confirmation of the transfer was performed by singleplex PCR method.

### 7. Data analysis

All data were entered into Microsoft Excel® and checked for completeness and consistency. The cleaned database was then imported into R software version 4.5.3 for statistical analysis. Categorical variables were summarized using frequencies and percentages. Comparisons between groups were performed using the chi-square test or Fisher’s exact test, as appropriate. Statistical significance was set at a p-value < 0.05

### 8. Quality control

Quality control of antibiotic discs was performed using *E. coli* ATCC 25922, *K. pneumoniae* ATCC 700603 (for *bla*_SHV_), *E. coli* ATCC 35218 (for *bla*_TEM_) and previously whole-genome sequenced *Enterobacterales* harboring all β-lactam and quinolone resistance genes found (unpublished data) were used as positive controls for PCR. *E. coli* ATCC45266 strains susceptible to rifampicin were used as control strain for plasmid transfer.

## RESULTS

### 1. Sociodemographic characteristics of the participants

A total of 754 consecutive clinical samples from patients who attended health facilities were processed in the microbiological laboratories of the YCH (59.5%; 449/754) and YUTH (40.4%; 305/754). Most samples originated from females 57.6% (434/754) and where aged >51 years old (34.4%; 259/754) (Figure 1). More than 60% of included patients were in-patients (Fig. 1).

**Figure 1:**
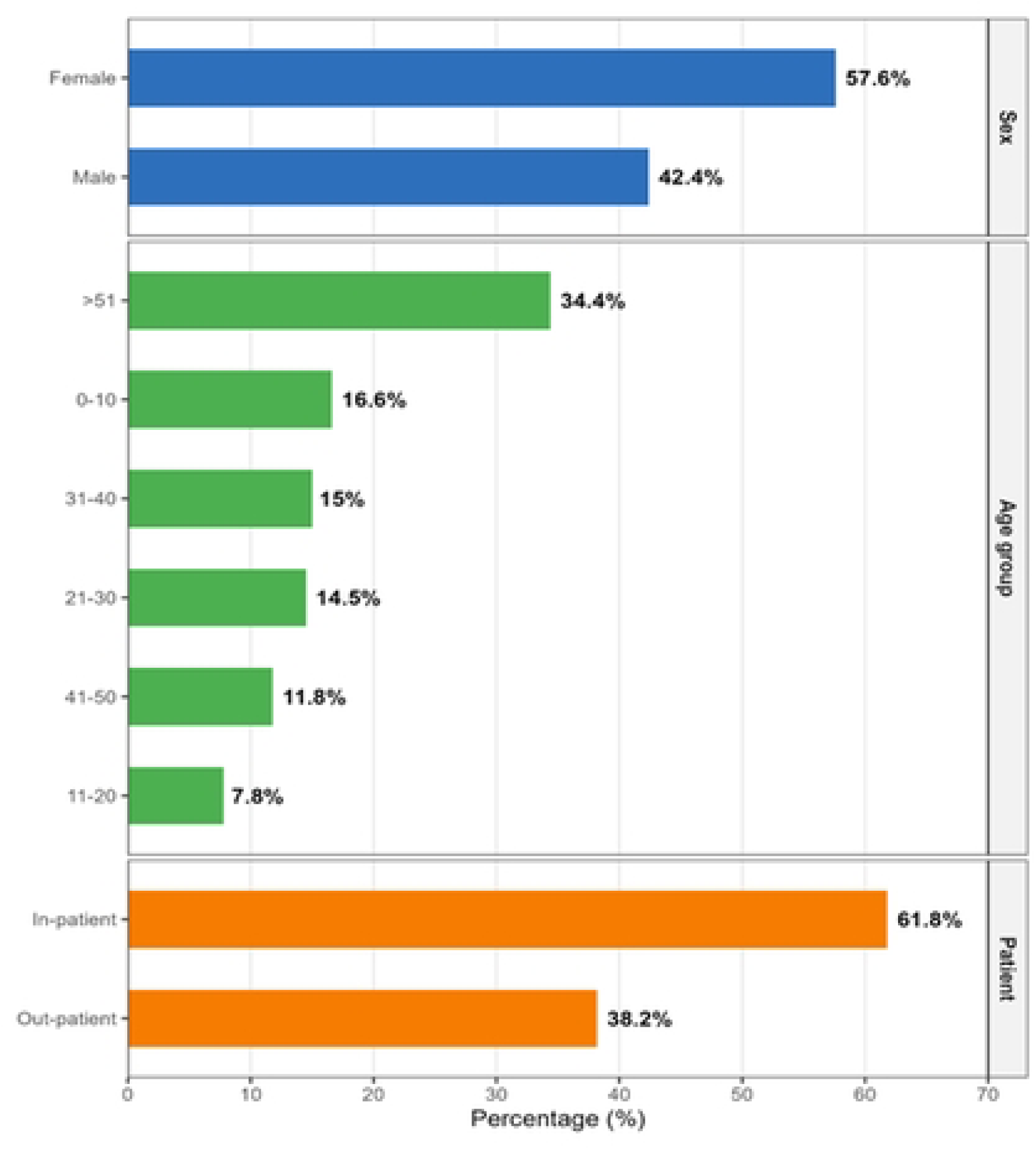
Distribution of patients according to sociodemographic characters

### 2. Prevalence and distribution of MDR, ESBL-producing and ciprofloxacin-resistant *Escherichia coli*

Out of 754 clinical specimens cultured, 24,9% (188/754) were positive with a total of 305 bacterial isolates being identified, of which *Klebsiella pneumoniae (*42.6%; 130/305) and *Escherichia coli* (21.3%; 65/305) were the predominant species of *Enterobacteriaceae.* Regarding the distribution of *E. coli* by specimen type, urine samples yielded the highest proportion of isolates (37,0%; 24/65), followed by stool (32,3%; 21/65) and pus (12,3%; 8/65). Of the 65 *E. coli* isolates, 50,7% (33/65) were classified as multidrug-resistant (MDR). Further characterization of the 33 *E. coli* isolates subjected to phenotypic ESBL testing revealed that all (100%) were ESBL producers, and 91% (30/33) exhibited resistance to ciprofloxacin (Fig. 2).

**Figure 2:**
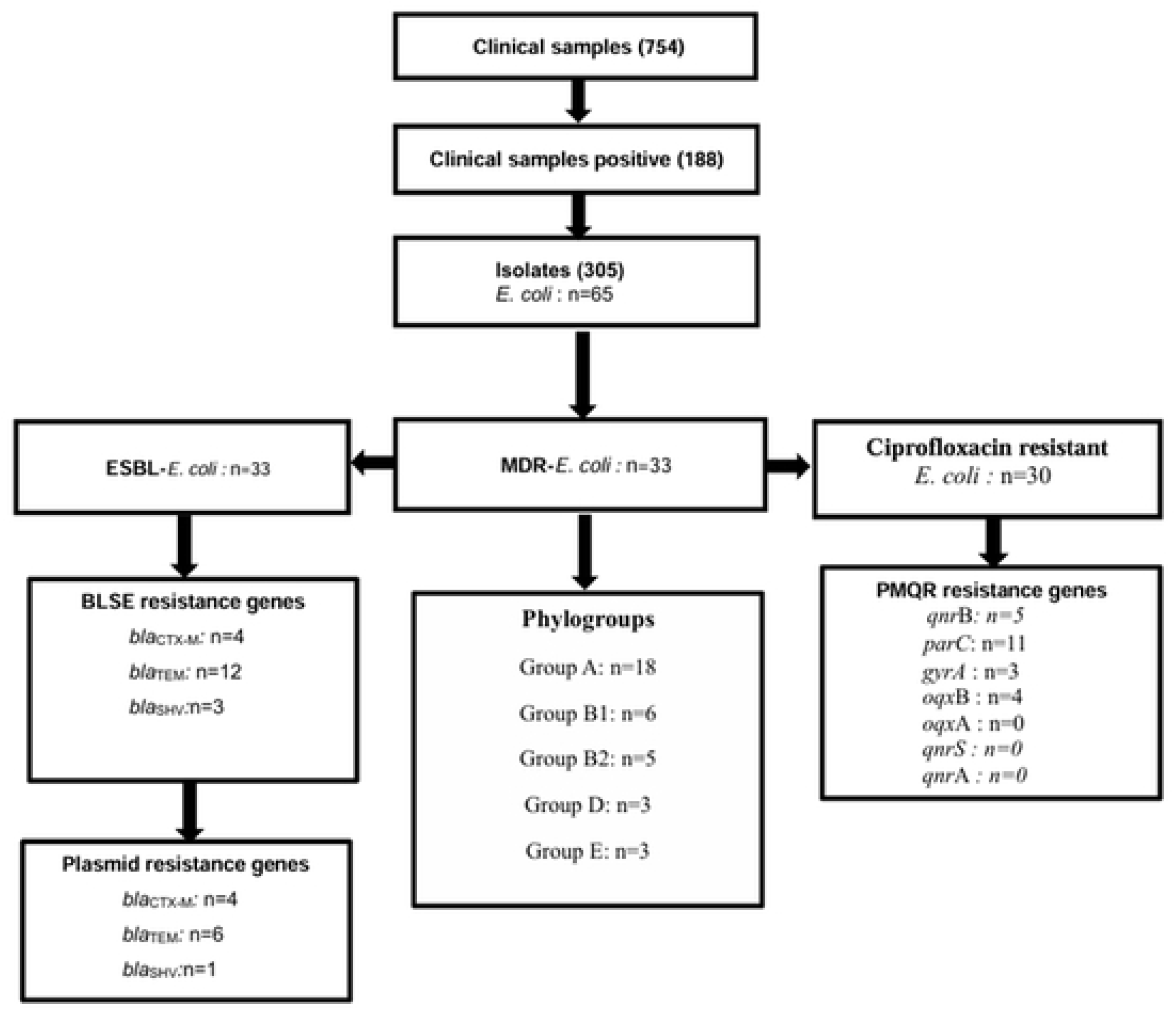
Diagram summarizing the results obtained

*MDR-E. coli* was more frequently recovered from in-patients (54.5%; 18/33) than from out-patients (45.5%; 15/33). Age-stratified analysis revealed that the highest number of *MDR*-*E. coli* isolates was recorded in patient aged over 51years, while the lowest was observed in the 0-10 years group (Table 1). These isolates were mainly from urine and stool samples (36.4%; 12/33) (Fig. 3).

**Figure 3:**
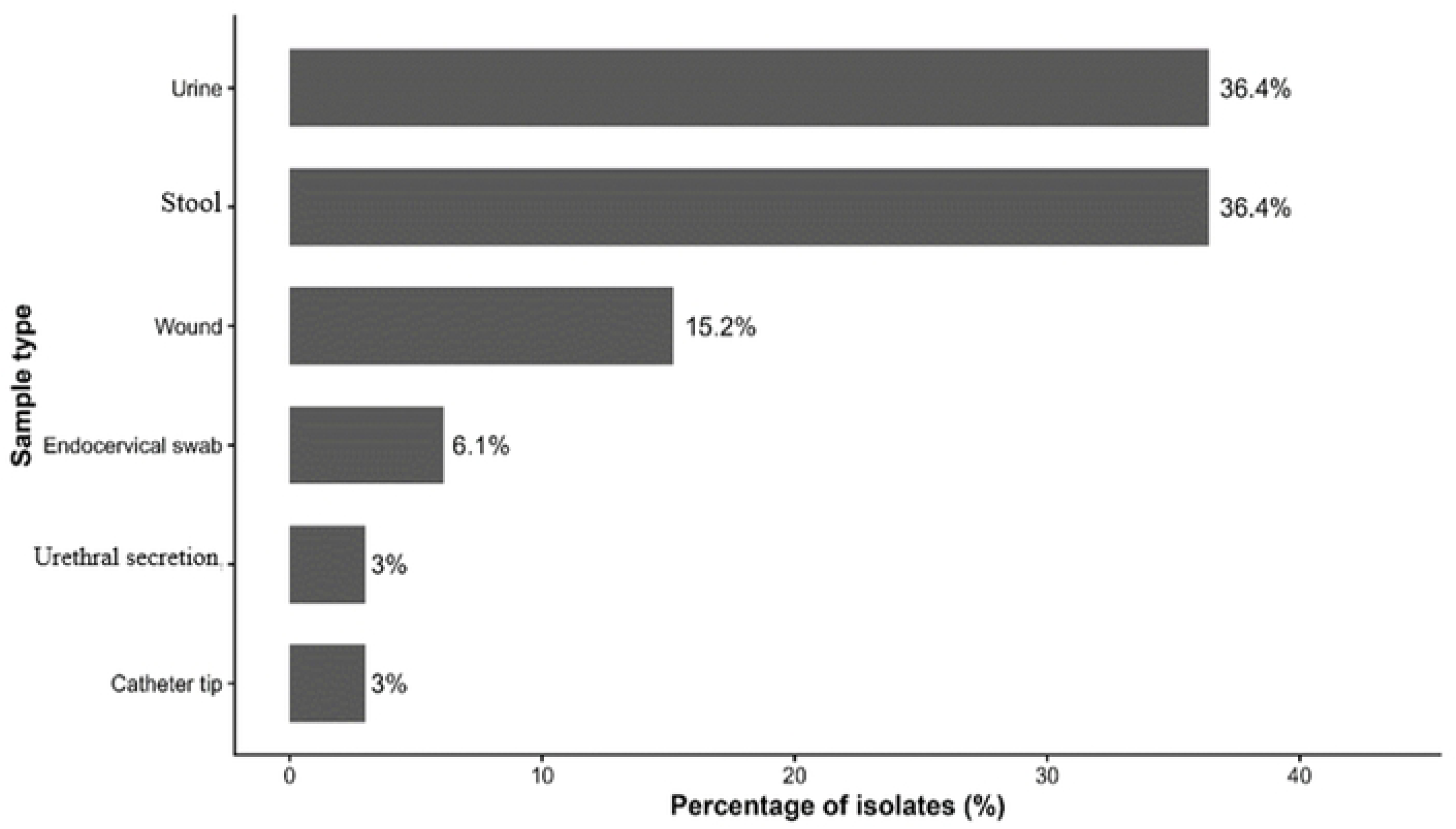
Distribution of *MDR Escherichia coli* isolates according to sample

**Table 1:**
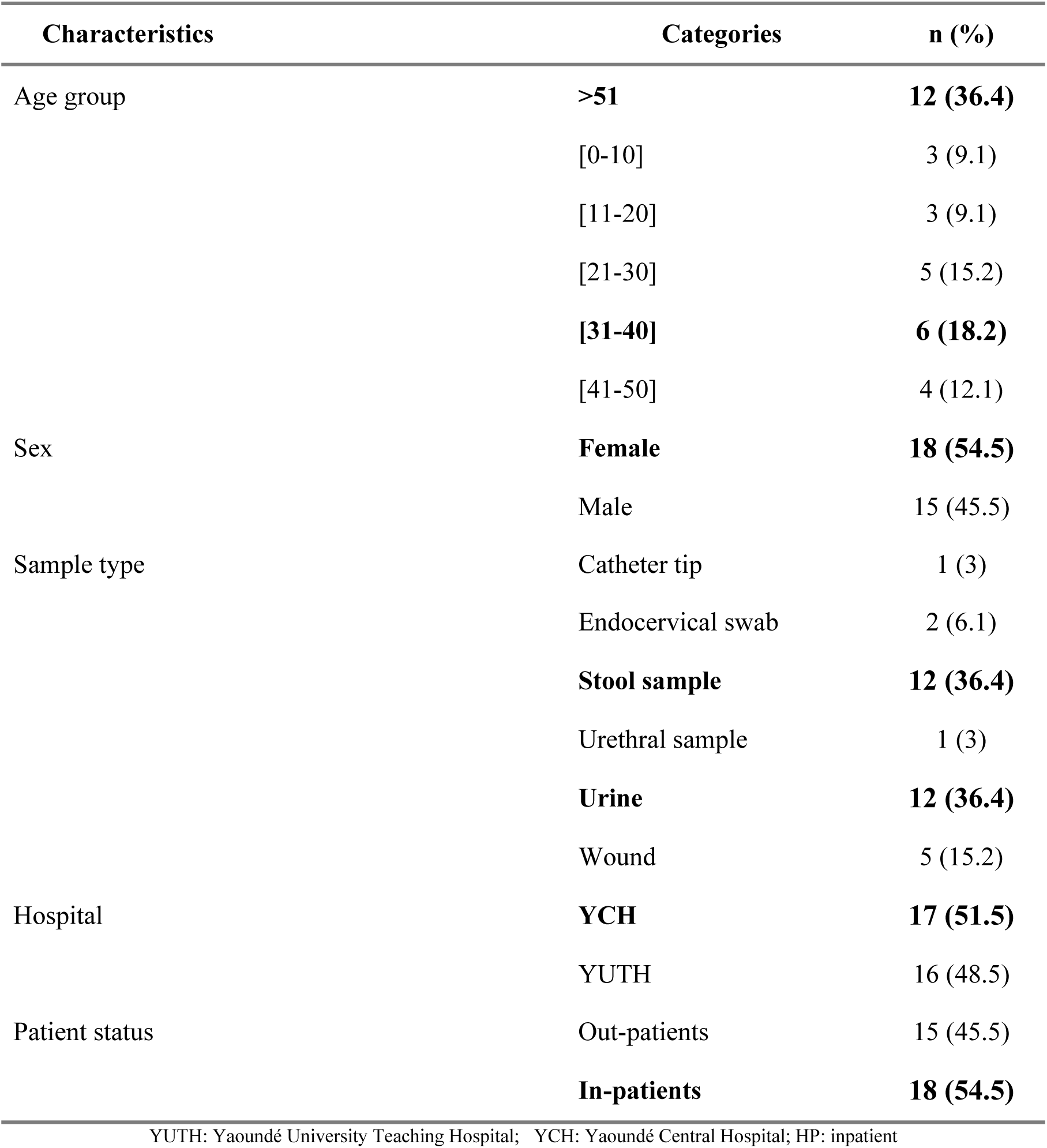
Distribution of multidrug-resistant *Escherichia coli* isolates according to sociodemographic and clinical characteristics

### 3. Antibiotic resistance profiles of MDR-*E. coli*

Multidrug-resistant *E. coli* has a high level of resistance to ceftriaxone (33/33; 100%), cefotaxime (33/33; 100%), followed by ciprofloxacin (30/33; 91%), norfloxacin (29/33; 88%), levofloxacin (28/33; 85%) gentamicin (23/33; 69%) and amoxicillin+ clavulanic acid (16/33; 48%). Meropenem and chloramphenicol were more effective with a low resistance rate (2/33; 6.1%), and (8/33; 24.2%) respectively (**Table 2**). The most prevalent resistant pattern was CRO-CAZ-CTX-CIP-CN-LEV-NOR (9/33; 28.1%) including seven antibiotics from three different families (**Fig. 4).**

**Figure 4:**
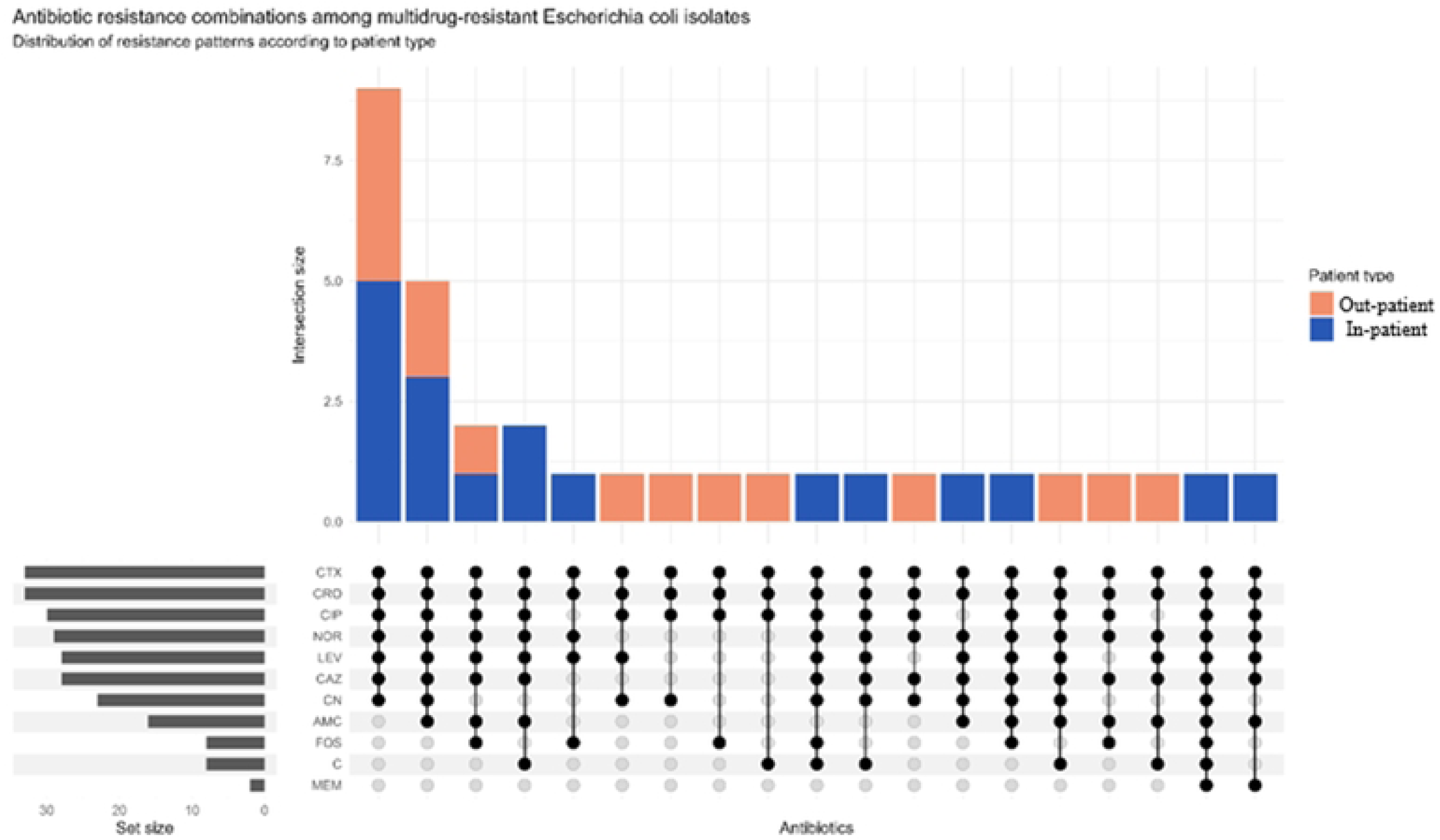
Antibiotic resistance combinations among multidrug-resistant *Escherichia coli*

**Table 2:** Antibiotic resistance profile

| Variable | Category | AMC | CRO | CTX | CAZ | MEM | CN | CIP | C | LEV | NOR | FOS |
| --- | --- | --- | --- | --- | --- | --- | --- | --- | --- | --- | --- | --- |
| Total isolates | MDR-E. coli (n=33) | 16 (48.5) | **33 (100)** | **33 (100)** | 28 (84.8) | 2 (6.1) | 23 (69.7) | **30 (90.9)** | 8 (24.2) | 28 (84.8) | **29 (87.9)** | 8 (24.2) |
| Hospital | YCH (n=17) | 6 (35.3) | **17 (100)** | **17 (100)** | 15 (88.2) | 2 (11.8) | 13 (76.5) | **16 (94.1)** | 5 (29.4) | 15 (88.2) | 15 (88.2) | 3 (17.6) |
|  | YUHC (n=16) | 10 (62.5) | **16 (100)** | **16 (100)** | 13 (81.2) | 0 (0) | 10 (62.5) | **14 (87.5)** | 3 (18.8) | 13 (81.2) | 14 (87.5) | 5 (31.2) |
|  | *p-value* | *0.169* | — | — | *0.656* | *0.485* | *0.465* | *0.601* | *0.688* | *0.656* | *1.000* | *0.438* |
| Patient type | In-patient (n=18) | 10 (55.6) | 18 (100) | 18 (100) | 17 (94.4) | 2 (11.1) | 13 (72.2) | 16 (88.9) | 5 (27.8) | 18 (100) | 18 (100) | 5 (27.8) |
|  | Out-patient (n=15) | 6 (40) | 15 (100) | 15 (100) | 11 (73.3) | 0 (0) | 10 (66.7) | 14 (93.3) | 3 (20) | 10 (66.7) | 11 (73.3) | 3 (20) |
|  | *p-value* | *0.491* | — | — | *0.152* | *0.489* | *1.000* | *1.000* | *0.699* | *0.013* | *0.033* | *0.699* |

YUHC: Yaoundé University Hospital; YCH: Yaoundé Central Hospital; HP: inpatient; Non-HP: non-hospitalized patient; AUG: amoxicillin + clavulanic acid, CAZ/cetazidime, CRO: ceftriaxone; CTX:

The comparison between the two hospitals revealed no statistically significant differences for all antibiotics tested (p > 0.05). In contrast, stratification by patient type showed significant differences for two fluoroquinolones: isolates from inpatients had significantly higher rates of resistance to levofloxacin (100% vs. 66.7%; p=0.013) and norfloxacin (100% vs. 73.3%; p=0.033) compared to those from outpatients, while no significant differences were observed for the other antibiotics tested (**Table 2).**

### 4. Prevalence of ESBL, PMQR resistance genes and mutation genes conferring the resistance of fluoroquinolones

The most prevalent ß-lactamase gene was *bla*_TEM_ (12/33; 36.4%) followed by *bla*_CTX-M_, (4/33; 12.1%) and *bla*_SHV_ (3/33; 9.1%) (**Table 3**). The most prevalent plasmid-mediated fluoroquinolone gene was *qnrB* (15.2%; 5/30) (**Table 3**).

**Table 3:** Prevalence of chromosomal ESBL, RQMP and mutation genes among multidrug-resistant *Escherichia coli* isolates

| Isolates n (%) | Chromosomal ESBL genes n (%) |  |  |
| --- | --- | --- | --- |
|  | *bla*<sub>CTX-M</sub> | *bla*<sub>TEM</sub> | *bla*<sub>SHV</sub> |
| *ESBL-MDR-E. coli*<br>(n=33) | 4 (12.1) | 12 (36.4) | 3 (9.1) |
| *In-patient*(18) | 4(22.2) | **7(38.8)** | 3(16.7) |
| *Out-patient*(15) | 0(0) | **5(33,33)** | 0(0) |

| Chromosomal RQMP and mutation genes n (%) |  |  |  |  |  |  |  |
| --- | --- | --- | --- | --- | --- | --- | --- |
| Isolates n (%) | *qnrA* | *qnrB* | *qnrS* | *oqxA* | *oqxB* | *parC* | *gyrA* |
| *ESBL-MDR-E. coli*<br>(n=30) | 0 (0) | 5 (16.6) | 0 (0) | 0 (0) | 4 (13.3) | 11 (36.6) | 3 (10) |
| *In-patient (15)* | 0 (0) | 2(13.3) | 0 (0) | 0 (0) | 1(6.6) | 8(53.3) | 2(13.3) |
| *Out-patient (15)* | 0 (0) | **3(20)** | 0 (0) | 0 (0) | **3(20)** | **3(20)** | 1(6.6) |

Among the 18 isolates from in-patients, 44.4% (8/18) carried at least one ESBL encoding genes, with *bla*_TEM_ the most prevalent, being detected in 38.8% (7/18) of isolates, followed by *bla*_CTX-M_ in 22.2% (4/18). Similarly, of the 15 isolates from out-patients, 46.7% (7/15) harboured at least one ESBL gene, with *bla*_TEM_ again the most frequently detected, accounting for 33.3% (5/15) of isolates **(Table 3).**

Regarding fluoroquinolone resistance, the analysis was performed on 30 ciprofloxacin-resistant isolates, of which 15 (50%; 15/30) were recovered from in-patient and 15(50 %; 15/30) from out-patient. Among the 15 isolates from in-patient, 20% (3/15) carried at least one PMQR gene, with *qnrB* the most prevalent, identified in 13.3% (2/15) of isolates. Whereas 26.6% (4/15) of the out-patient harboured at least one PMQR gene, with oqxB and qnrB the most frequently detected, each found in 20% (3/15) of isolates **(Table 3).**

Molecular analysis of these ciprofloxacin-resistant isolates revealed that chromosomal mutations were the predominant resistance mechanism conferring fluoroquinolone resistance. Specifically, the *parC* gene mutation was the most frequently detected, with 36.6% (11/30) of isolates carrying the Serine (Ser)80→ isoleucine (Ile) substitution, followed by Serine (Ser)83→ isoleucine (Ile) substitution in the *gyrA* gene (10%; 3/30) **(Table 3).**

### 5. Prevalence of plasmid-mediated ESBL and PMQR resistance genes

Among the 15 ESBL-producing isolates carrying at least on plasmid-mediated resistances gene, *bla*_TEM_ was the most frequently detected, identified in 80% (12/15) of isolates, followed by *bla*_CTX-M_ in 26.6% (4/15). Furthermore, of the 12 isolates carrying *bla*_TEM_ and the six isolates carying *bla*_CTX-M_, 50% (06/12) and 100% (4/4) of these genes, respectively were located on plasmids (Fig. 5).

**Figure 5:**
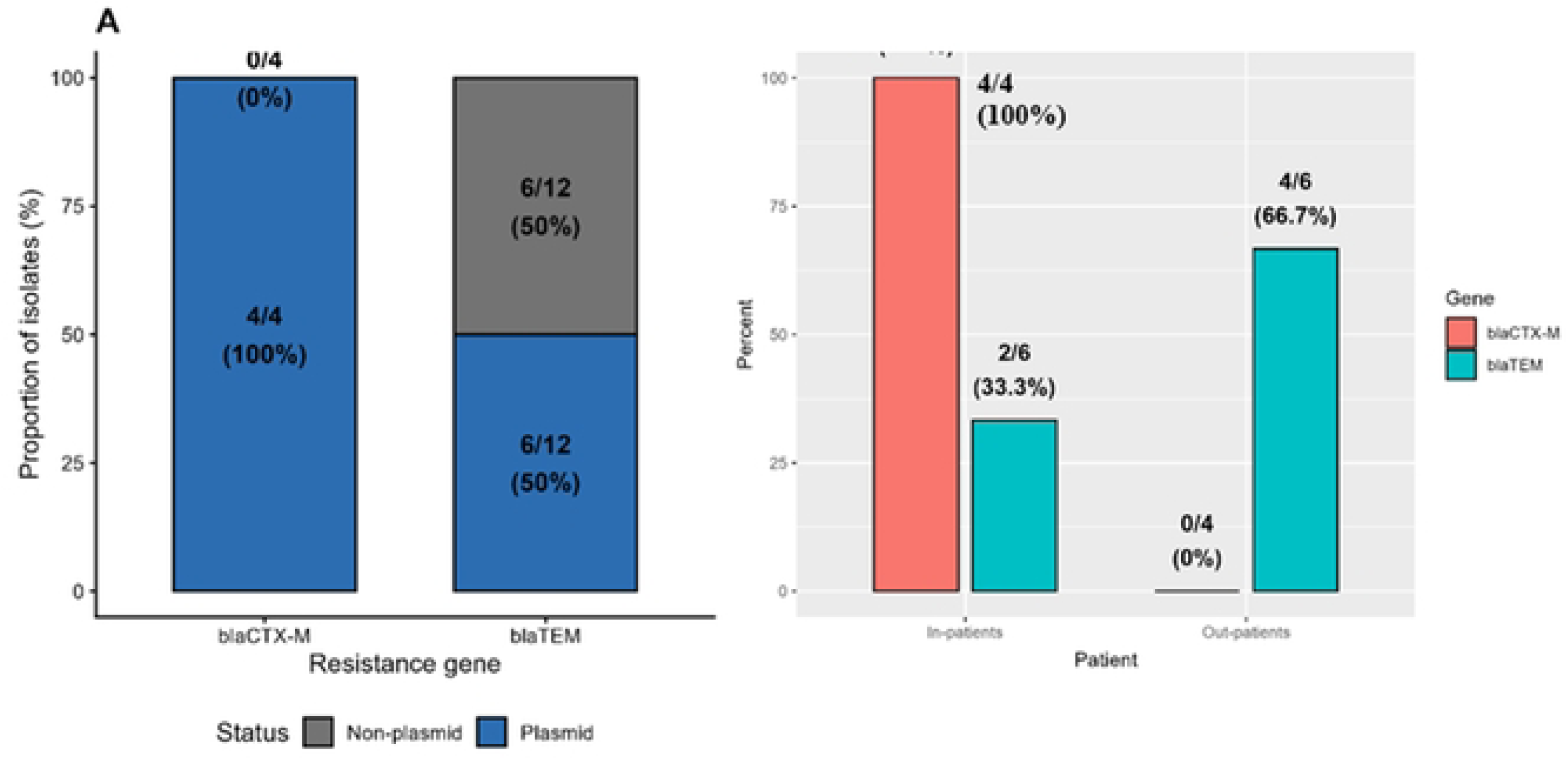
Distribution of plasmid-mediated ESBL genes among ESBL-producing isolates

The *bla*_CTX-M_ predominated among in-patients (100%; 4/4) while the *bla*_TEM_ was the most prevalent plasmid resistance gene in out-patients (66.6%; 4**/**6) (Fig. 5). Nevertheless, despite the plasmid-borne location of these resistance genes, conjugation experimentation revealed no horizontal gene transfer to recipient strains (Fig. 5).

### 6. Phylogroup distribution

Phylogroup analysis of the 33 *E. coli* multidrug-resistant isolates revealed the presence of four phylogenetic groups (A, B1, B2, D). Phylogroup A was the most predominant (54.5%; 18/33) followed by phylogroups B1 (18.2%; 6/33); B2 (15.2%; 5/33); and phylogroup D (9.1%, 3/33). phylogroup B2 which is associated with virulent *E. coli* was the most frequently detected in stool (80%; 4/5) and urine sample (20%; 2/5) (Fig. 6).

**Figure 6:**
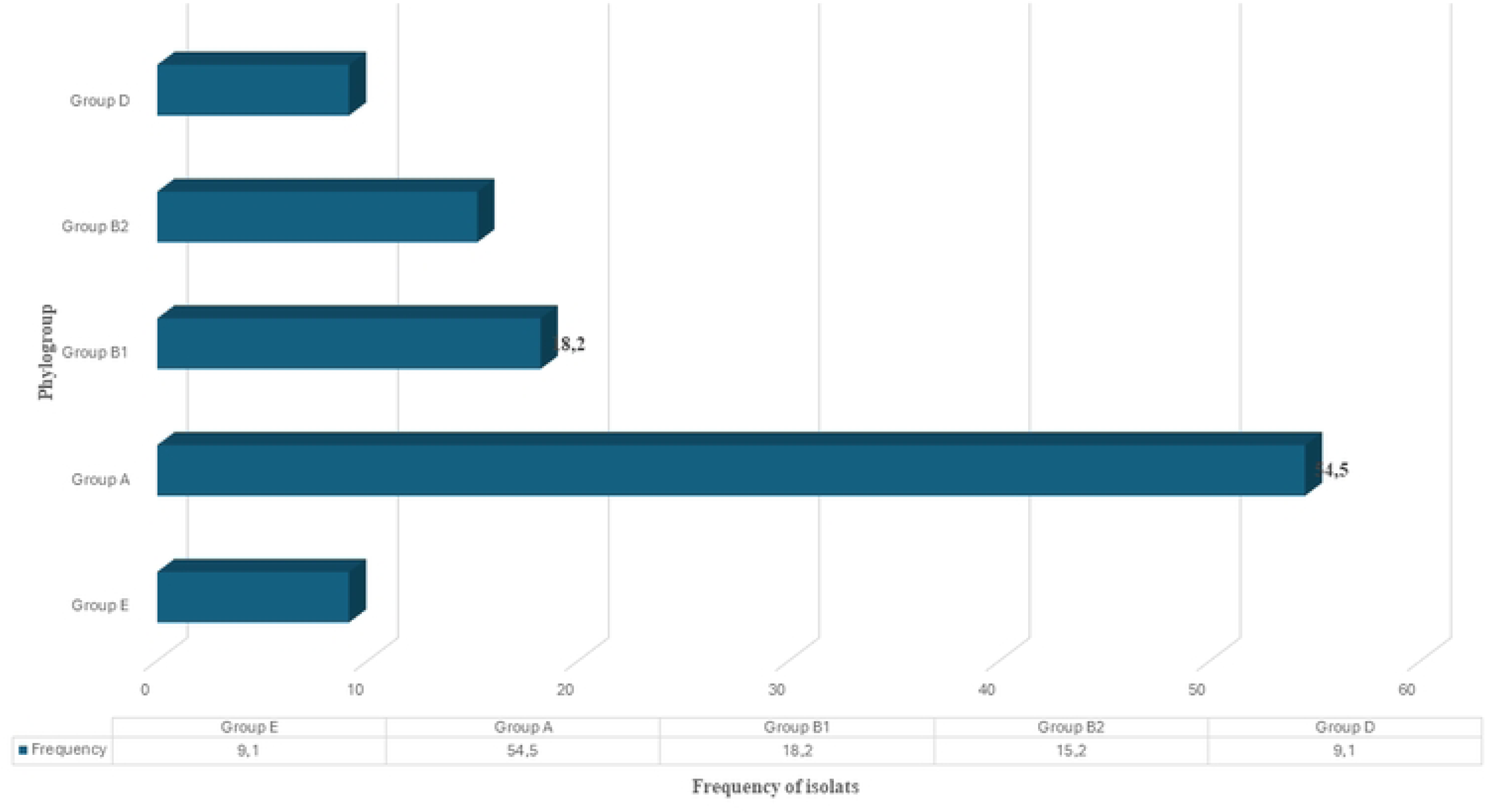
Frequency of phylogroup

## DISCUSSION

Since the discovery of antibiotics, their misuse has contributed to the emergence and escalation of AMR, of which a significant proportion is driven by *ESBL-Ec* and *MDR-Ec* (25). The purpose of this study was to investigate investigate the prevalence, phenotypic profiles and genetic characteristics including, plasmid borne antibiotic resistance genes and point mutation regions conferring resistance to fluoroquinolones among *MDR-Ec isolated* from clinical samples in two hospitals in Yaounde, Cameroon.

Overall, 50,7% (33/65) of *E. coli* were multidrug-resistant. This finding aligns with data generated in the West region of Cameroon, by Bayaba et al. (2025) who reported a high prevalence of 85.71% of MDR *E. coli* among urinary isolates (6). However, our findings are lower than that reported by the surveillance conducted at the National Reference Hospital in Cameroon (Centre Pasteur du Cameroun, Yaoundé), documented an overall antibacterial resistance rate of 96.0% among priority pathogens between 2010 and 2017, with a marked rise in third-generation cephalosporin resistance among *Enterobacteriaceae* over this (26). Despite these differences in prevalence, these findings remain no less alarming, as even a relatively low resistance rate constitutes, in itself, a major public health concern. All MDR isolates identified in this study were concomitantly ESBL-producers (MDR-ESBL), and the absolute resistance observed to ceftriaxone and cefotaxime among these MDR-ESBL isolates is particularly alarming. This finding reflects the strong selective pressure exerted by third-generation cephalosporins, which are among the first-line antibiotics prescribed in health facilities and available over the counter in Cameroon (27, 28). This generalized resistance of C3G is consistent with the high prevalences of the ESBL-coding genes identified in this study, as the presence of these enzymes hydrolyzes ß-lactam cycles of C3G, making their clinical efficacy limited (27). Of the multidrug-resistant-ESBL-*E. coli* isolates, 45.5% carried at least one gene encoding an ESBL. The *bla*_TEM_ gene was the most prevalent (36.4%), followed by the *bla*_CTX-M_ gene (12.1%). These results diverge from those reported by Ouchar Mahamat et al. (2021) in Chad and Bayaba et al. (2025) in Cameroon, who identified *bla*_TEM_ as the dominant ESBL gene among clinical *E. coli* isolates from urine samples (6, 29).

Resistance to fluoroquinolones is also alarming with 91% of MDR-*Ec* isolates resistant to ciprofloxacin and 88% to norfloxacin. This resistance is of great concern, given that this family of antibiotics is commonly prescribed in Cameroon as empirical treatment, especially for urinary tract infections, typhoid fever, urogenital infections and severe infections such as sepsis Their extensive use, combined with inadequate antimicrobial stewardship, weak enforcement of regulations may contribute to the emergence and persistence of fluoroquinolone resistance (26, 30). Two distinct resistance mechanisms co-occurred among these isolates: chromosomal QRDR mutations (*gyrA*, *parC*) and the plasmid-mediated *qnrB* gene, which together drastically compromises the therapeutic efficacy of this class of antibiotic (31). The *qnrB* gene was identified in 16.6% of isolates, this is prevalence is higher than that reported by Lyonga et al. (2020) in Yaoundé, where the *qnrB* gene was detected in only 4.4% of *E.coli* isolates isolated of clinical sample at the Yaounde University Teaching Hospital, Yaounde General Hospital and the Yaoundé Gynaeco-Obstetric and Pediatric Hospital between 2013 and 2015 (10), this resistance could be explained by the fact that, faced with the emergence of beta-lactam-resistant and beta-lactamase-producing bacteria, therapeutic approaches had to be adapted, making quinolones a class of antibiotics now administered in Cameroon.(6, 32). Several structural challenges are causing this increase in antibiotic resistance, particularly in Cameroon, where self-medication and over-the-counter antibiotics create favourable conditions for the emergence of multi-resistant bacteria. These results underscore the urgent need to revise empirical treatment protocols in Cameroon, strengthen antibiotic stewardship across health facilities, and implement strategies to curb the over-the-counter sale of antibiotics without prior prescription. Although Cameroon adopted a National Action Plan (NAP) on antimicrobial resistance, its implementation has remained limited to the human sector, and the country still lacks a dedicated, functional AMR containment programme. Addressing these structural gaps is essential to translate policy commitments into effective, sustained control of AMR spread(33). Despite the high prevalence of resistance, meropenem remained active against the majority of *E. coli* isolates, with a resistance rate of 6.1%. This result is consistent with the study by Koudoum et al. (2025), in which meropenem was the most active antibiotic against ESBL-producing *E. coli* isolates; nevertheless, these isolates obtained from clinical, environmental, and hospital wastewater samples exhibited a meropenem resistance rate of 21.33% (16). These findings underscore the need to preserve carbapenems for severe *E. coli*-related infections, in line with their classification as Reserve-group agents under the WHO AWaRe framework, which restricts their use to confirmed multidrug-resistant infections in order to safeguard their long-term efficacy(*34, 35*). However, even the minimal emergence of resistance to carbapenems is of concern and requires continued monitoring, given that carbapenem-resistant *Enterobacteriaceae* are associated with extremely high levels of treatment options.

Plasmid analysis revealed that 66.6% of isolates carrying ESBL genes hosted these genes on their plasmid DNA, with *bla*_TEM_ detected on plasmids in 42.85% of cases and *bla*_CTX-M_ in 28.5%. This high proportion of ESBL genes carried by plasmids is consistent with the well-known ability of these determinants to be mobilized on conjugative or mobilizable plasmids, facilitating their horizontal transfer between strains and species (36, 37). A comparable distribution was reported by in Nepal, where *bla*_TEM_ and *bla*_CTX-M_ were detected on plasmids in 63.4% and 36.6% of isolates respectively, reinforcing the global importance of plasmid dissemination of the ESBL genes (38).

In contrast, no PMQR genes were detected on plasmid DNA in this study, suggesting that in the isolates examined, fluoroquinolone resistance is primarily maintained by chromosomal mechanisms namely *gyrA* and *parC* mutations rather than by plasmid-mediated determinants. Although genes that are classically described as carried by plasmids and often associated with integrons and other mobile genetic elements (39) several studies have documented rare chromosomal integration of PMQR genes through transposition events, particularly in isolates subjected to prolonged antibiotic pressure (31). The absence of plasmid-based PMQR determinants suggests a chromosomal stabilization of this resistance, probably favored by sustained antibiotic pressure. In fact, the selective pressure exerted by the use of fluoroquinolones could promote the chromosomal binding of *qnr genes*, a phenomenon that confers greater stability to resistance and reduces the likelihood of gene loss in the absence of antibiotic pressure (40). This hypothesis should be deepened by whole genome sequencing approaches in order to precisely characterize the genomic context of these resistance determinants.

Phylogroup A was the most prevalent among MDR-*Ec* isolates, followed by group B1. These two groups are classically associated with commensal strains of *E. coli* (41). The predominance of multidrug-resistant among commensal phylogroups is of great concern because these are natural gut colonizers and therefore represent major reservoirs of resistance genes. They also constitute a critical interface for horizontal gene transfer to pathogenic strains by conjugation, transformation or transduction (41, 42). In the Cameroonian context where faecal contamination of water sources and food is a persistent challenge, the widespread circulation of commensal and multidrug-resistant *E. coli* in the community represents an important reservoir and distributor of resistance, which could accelerate the spread of AMR determinants to clinically dangerous pathogens (43) Strengthening surveillance systems integrating both clinical and community isolates in antimicrobial resistance in Cameroon.

This study has some limitations. First, the analysis of ESBL and PMQR gene distribution was based on relatively small subgroup sizes which limits the precision of the reported prevalence estimates and the statistical robustness of between-group comparisons. Second, the study was conducted in two hospitals in Yaoundé, and the findings may therefore not be generalizable to rural health facilities, primary care settings, or other regions of Cameroon, where antibiotic prescribing practices and selective pressures may differ. Third, although this work is framed within a One Health perspective, the isolates analyzed were exclusively of clinical origin; the absence of parallel environmental or animal sampling limits our ability to empirically capture the cross-sectoral transmission pathways discussed. Finally, resistance gene detection relied on targeted PCR rather than whole-genome sequencing; this approach does not allow confirmation of the exact allelic variants detected (e.g., distinguishing non-ESBL-conferring *bla_TEM-1_*_/TEM-2_ from ESBL-conferring TEM variants), and may therefore underestimate the true genetic diversity of the resistance determinants circulating in this setting.

## CONCLUSION

This study reveals an alarming prevalence of MDR-*Ec* isolated from two health facilities in Yaoundé, with all being ESBL-producers (dominated by the *bla_TEM_*), and almost widespread resistance to fluoroquinolones. These results call for an urgent strengthening of antibiotic stewardship policies in Cameroon, a review of empirical treatment protocols, and the establishment of routine, real-time One Health genomics surveillance systems. Integrated One Health surveillance, involving the clinical, community and environmental sectors, is equally essential to limit the spread of antimicrobial resistance in this resource-limited context.

## Abbreviations

AMR: antimicrobial resistance
MDR: Multidrug resistance
ESBL: Extended Spectrum β-lactamase
PMQR: Plasmid-mediated Quinolone Resistance
PCR: Polymerase Chain Reaction
WHO: World Health Organization

## Author contributions

**PKM**: Sample and data collection, bacteria culture and identification, molecular analysis, data interpretation and manuscript drafting. **LLF**: critically revised the manuscript **AAZ:** Sample and data collection, bacteria culture and identification, molecular analysis. **JVM**: bacteria culture and identification, molecular analysis **JKN**: critically revised the manuscript. **HG**: contributed to the study design and facilitated the implementation of the study. **RCF:** Supervised the study and critically revised the manuscript

## Acknowledgements

The authors sincerely thank all the study participants and the members of the CEDBCAM-RI group for their valuable support and contributions throughout this study

## List of supplementary table and figure

**Table sup I:**
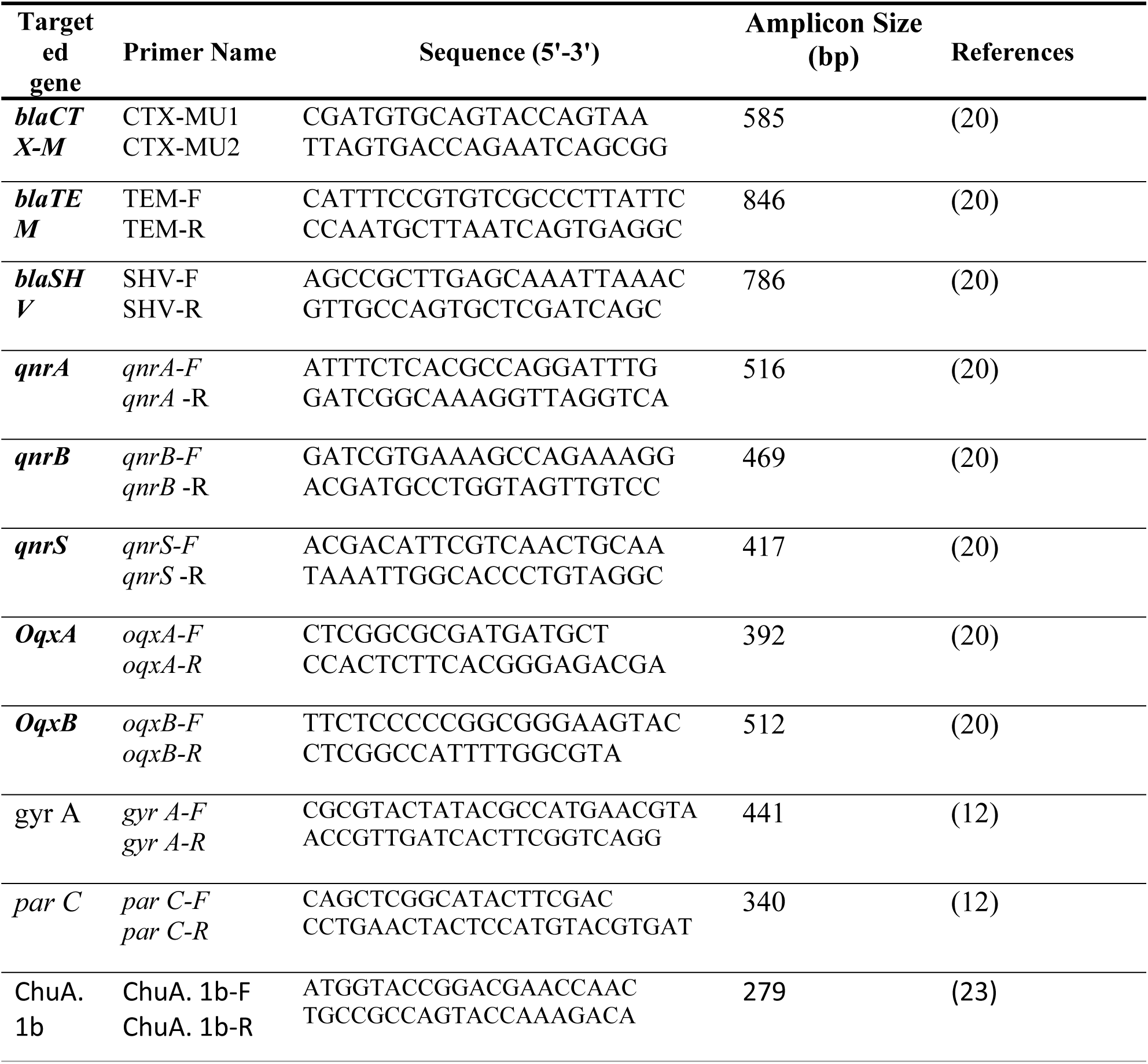
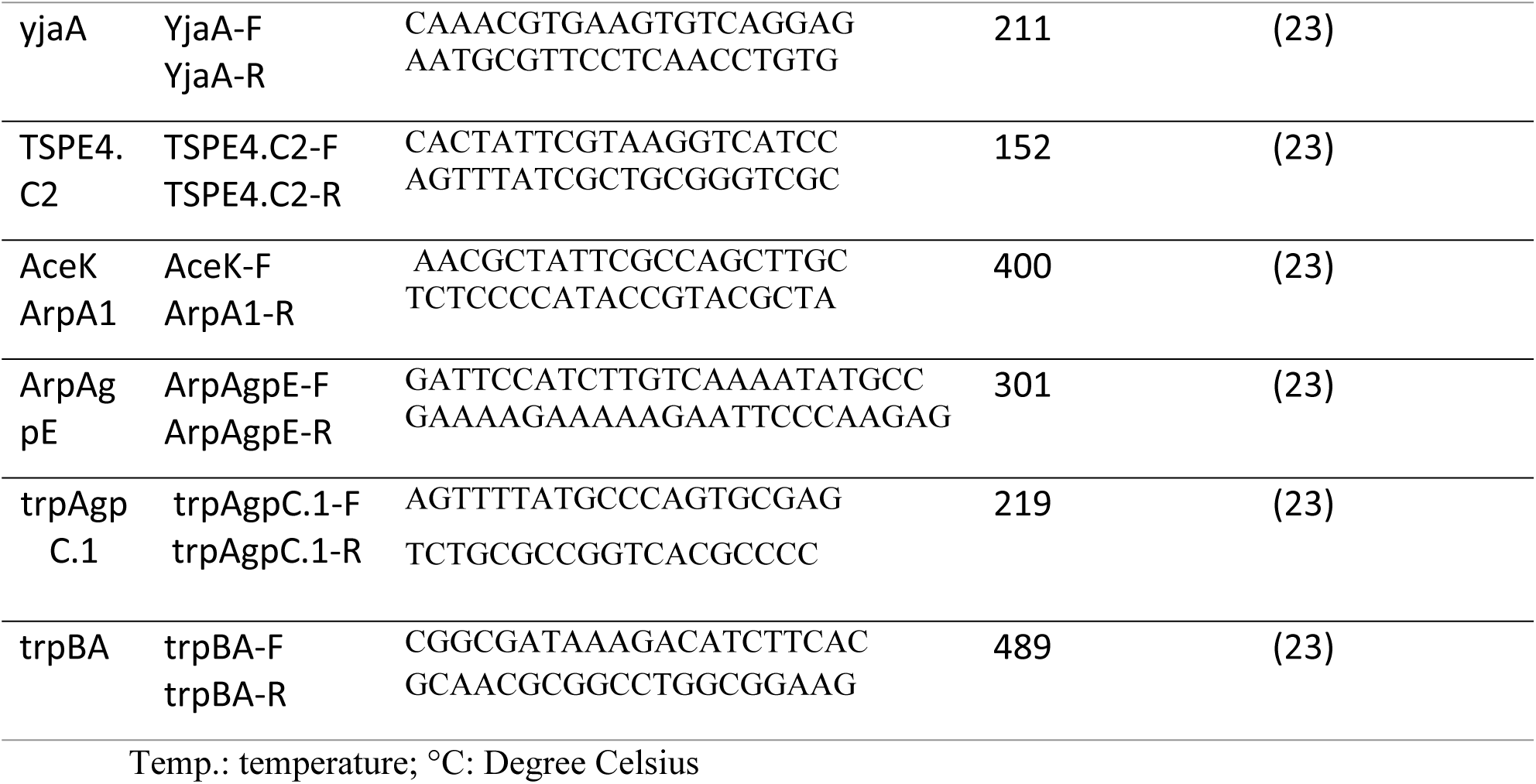
List of primers used for the amplification of the PMQR and ESBL genes.

